# Reputation risk during dishonest social decision-making modulates anterior insular and cingulate cortex activity and connectivity

**DOI:** 10.1101/2022.11.28.518136

**Authors:** Lennie Dupont, Valerio Santangelo, Ruben Azevedo, Maria Serena Panasiti, Salvatore Maria Aglioti

**Affiliations:** Department of Psychology, Sapienza University of Rome, Via dei Marsi 78, Rome, Italy; IRCCS Fondazione Santa Lucia, Via Ardeatina 306, 00179 Rome, Italy; Department of Philosophy, Social Sciences & Education, University of Perugia, Piazza G. Ermini 1, 06123 Perugia, Italy; Keynes College, School of Psychology, University of Kent, Canterbury, Kent, CT2 7NP, United Kingdom; Sapienza University of Rome and CLN^2^S@Sapienza, Italian Institute of Technology, Rome, Italy

**Keywords:** Deception, decision-making, lie, morality, reputation risk, fMRI

## Abstract

To explore the neural underpinnings of (dis)honest decision making under quasi-ecological conditions, we used an fMRI adapted version of a card game in which deceptive or truthful decisions are made to an opponent, with or without the risk of getting caught by them. Dishonest decisions were associated to increased activity in a cortico-subcortical circuit including the bilateral anterior cingulate cortex (ACC), anterior insula (AI), left dorsolateral prefrontal cortex, supplementary motor area, and right caudate. Crucially, deceptive immoral decisions under reputation risk enhanced activity of – and functional connectivity between – the bilateral ACC and left AI, suggesting the need for heightened emotional processing and cognitive control when making immoral decisions under reputation risk. Tellingly, more manipulative individuals required less involvement of the ACC during risky self-gain lies but more involvement during other-gain truths, pointing to the need of cognitive control only when going against one’s own moral code.

## Introduction

Although harmful to interpersonal interactions in financial, political, legal and also daily-life contexts, dishonesty remains ubiquitous. Classically investigated by economic, sociological and psychological sciences, only in the last decades has dishonesty attracted the interest of the neurosciences. Specifically, an increasing number of functional neuroanatomy studies have tried to untangle the complex mechanisms behind dishonest decision-making ^1–17^. Activation Likelihood Estimation (ALE) meta-analyses on the neural correlates of deception vs. non-deceptive behavior ^18–20^ show the involvement of a large cortico-subcortical neural network associated with complex functions related to deception, such as perspective taking, emotional, moral, and reward processing. In sum, dishonesty seems to mainly recruit anterior portions of the brain, such as the dorsolateral prefrontal cortex (dlPFC), the ventromedial prefrontal cortex (vmPFC), the anterior insula (AI), the anterior cingulate cortex (ACC), and the nucleus caudate (Cau), although posterior activation of brain regions, such as the inferior parietal lobule (IPL) and temporoparietal junction (TPJ) has also been reported. It has been suggested that successful deceiving increases activity in areas related to executive/cognitive control possibly because of the need to inhibit predominant truthful responses^8,19^. However, given the complexity of deception, a variety of personality, cognitive and emotional factors seem to orchestrate controlled and automatic decision-making across individuals, with some being highly susceptible to the temptation to lie (Will hypothesis) and others being ‘naturally’ immune to moral violations (Grace hypothesis)^12^. Importantly, it seems that cognitive control may allow cheaters to behave honestly and honest people to cheat depending on the circumstance^21^.

It is worth noting that the initial exploration of the neural correlates of dishonest decision-making was based on tasks where the experimenter specifically instructed participants when to lie and when to tell the truth in impoverished laboratory conditions. Thus, lack of intentionality and absence of social contexts made dishonest decision-making very different from what happens in real-life conditions^22^. Aware of the need to explore dishonesty in improved contexts^23^, subsequent neuroimaging studies have used paradigms devised to circumvent the oversimplified laboratory conditions that are unfit to capture the complexity of (dis)honest decision-making^12,22,24^. Yet, most of the existing studies missed one or more of the features that make a laboratory paradigm as ecological as possible, namely, i) intention to lie, i.e., when the choice to lie is spontaneous instead of being instructed; ii) social interaction, i.e., when the deception occurs in a social context; and iii) motivation, i.e., when telling a lie entails a benefit or avoidance of a penalty for the liar^22^. To explore the pattern of neural activity during dishonesty in quasi-ecological conditions, we combined fMRI with a new version of a behavioral task, the temptation to lie card game (TLCG), that we developed in previous studies and that proved adept to tap multiple facets of spontaneous social deception^25–30^.

The TLCG is an interactive card game where experimental participants observe another player (who unbeknownst to the participant was a computerized opponent) choosing one out of two covered cards, namely, the ace of heart or of spade. Picking one or the other implied winning or losing money, respectively, from a common pocket. The participant was the only one who could see the choice outcome and had the liberty to accept or revert the outcome. For example, when the outcome implied a loss for the participant they could report a win instead, thus making a dishonest decision (self-gain lie). Therefore, the TLCG includes intentionality, by letting the participants make spontaneous decisions to lie or tell the truth, sociality, by including an opponent whom the participant played against, and motivation, by including a monetary reward when they win on a trial-by-trial basis.

An important and somewhat neglected facet of deceptive dishonesty explored by the our approach is the probability of getting caught in one’s lies. Indeed, our participants were informed that in some trials nobody could see their decision (no-reputation risk) while in others the opponent could be informed about their decision (reputation risk). Thus, any immoral decision jeopardized the participant’s moral reputation in the latter but not in the former type of trial. Of note, the loss of reputation is so relevant that people may prefer undergoing very unpleasant experiences, such as physical pain, to avoid it^31^. Tellingly, the risk of being caught while lying, not only decreases the tendency to deceive^26^ but it is also related to the need of regulating the activity of the sympathetic nervous system^25^ and to cardiac interoception^29^.

The main aim of our study was to determine the neural network involved in spontaneous dishonest decision-making in general and in the processing of lies, as well as its modulation by the risk of getting caught by the opponent and thus losing their reputation. Moreover, we explored whether individual differences in morality were associated with the modulation of dishonest decision contingent upon reputation risk.

## Results

### fMRI results

We used a first general linear model (GLM1; see section “fMRI data analysis”) to investigate the main effect of spontaneous lying vs. truth-telling and the interaction between spontaneous lying and the reputation conditions (reputation (Rep), vs. no reputation, (NoRep)), independently of the outcome (i.e., favorable (Fav) or unfavorable (UnFav)). Spontaneous lying (vs. truth-telling) recruited a brain circuit involving anterior regions, including the anterior cingulate cortex (ACC), the anterior insula (AI), bilaterally, the left dorsolateral prefrontal cortex (dlPFC), the right supplementary motor area (SMA), and the right caudate (Cau) (Fig. 1a, purple maps, 1b grey bar plots & Table 1). These regions showed a selective increase of activity for lie conditions irrespective of reputation condition (Fig. 1b: compare bars 1 & 3 vs. bars 2 & 4). A significant interaction effect of reputation (Rep, NoRep) x decision (Lie, Truth) was also found, driven by increased activity in the ventromedial portion of the left anterior insula for Rep_Lie (vmAI; yellow bar plot in Fig. 1b & Table 1). The related signal plot shows increased activity in this region when participants lied under reputation risk as compared to the other conditions (compare bar 3 vs. the other bars in the vmAI signal plot of Fig. 1b). No other interaction effects were found within this model.

**Table 1.** Brain activation for the main effect of “spontaneous lies vs. truths” (NoRep_Lie +Rep_Lie > NoRep_Truth + Rep_Truth) and for the interaction (Rep_Lie + NoRep_Truth > NoRep_Lie + Rep_Truth). Only clusters and their local maxima that survived family-wise error correction (p <.05) are presented.

|  | L/R | Regions | kE | x | y | Z | Z | P-FWE-corrected |
| --- | --- | --- | --- | --- | --- | --- | --- | --- |
| Main effect of Lie |  |  |  |  |  |  |  |  |
|  | R | ACC | 1835 | 4 | 28 | 28 | 5.69 | <0.001 |
|  | R | ACC |  | 4 | 36 | 28 | 5.66 | <0.001 |
|  | L | ACC |  | -6 | 28 | 28 | 5.32 | <0.001 |
|  | L | ACC |  | -10 | 30 | 20 | 5.13 | <0.001 |
|  | R | ACC |  | 6 | 44 | 20 | 4.75 | 0.002 |
|  | L | ACC |  | 0 | 48 | 18 | 4.7 | 0.002 |
|  | L | AI | 237 | -34 | 22 | -8 | 5.52 | <0.001 |
|  | L | AI |  | -42 | 16 | -8 | 5.01 | <0.001 |
|  | R | AI | 99 | 30 | 18 | -10 | 4.76 | 0.002 |
|  | R | SMA | 158 | 10 | 14 | 62 | 4.19 | 0.018 |
|  | L | dIPFC | 23 | -22 | 46 | 30 | 4.09 | 0.026 |
|  | R | CAU | 19 | 12 | 14 | 12 | 4.02 | 0.034 |
| Interaction: Rep - No Rep x Lie - Truth |  |  |  |  |  |  |  |  |
|  | L | AI | 60 | -32 | 22 | -10 | 4.04 | 0.031 |

Abbreviations: L/R, left/right; kE, number of voxels; p-FWE, family-wise error corrected p-value; ACC, anterior cingulate cortex; AI, anterior insula; SMA, supplementary motor area; dIPFC, dorsolateral prefrontal cortex; CAU, caudate.

**Figure 1.**
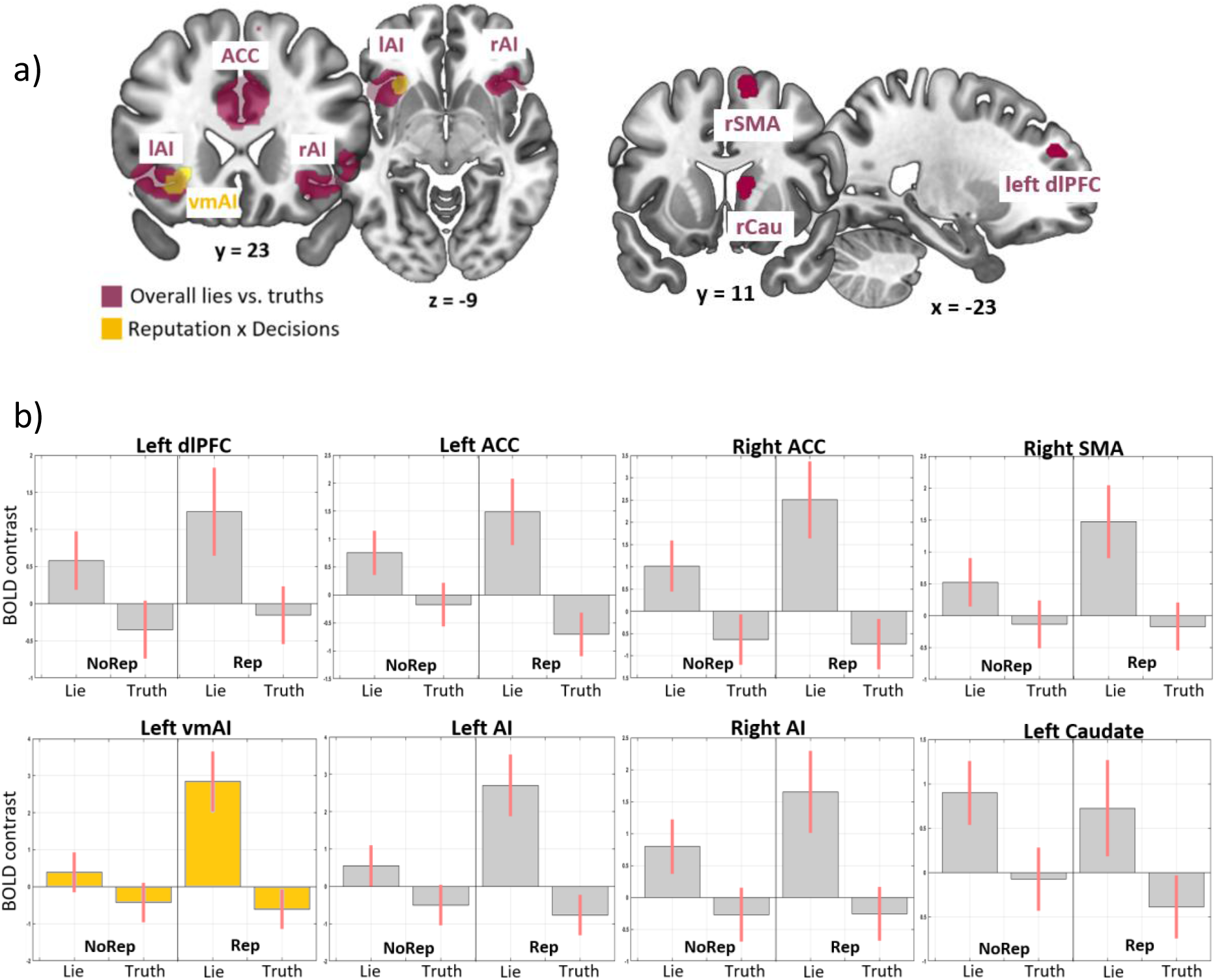
Results of GLM1: a) coronal and axial slices showing the regions active for lies vs. truths in both conditions, i.e., ACC and AI (purple map) and the left ventromedial section of the left AI for lies in the reputation risky condition (yellow map). b) Bar plots of regions more active for lies vs truths. Grey bar plots for lying in both conditions, yellow bar plots for spontaneous lies in the reputation risk condition. Abbreviations: ACC, anterior cingulate cortex; lAI/rAI, left/right anterior insula; rSMA, right supplementary motor area; dlPFC, dorsolateral prefrontal cortex; rCau, right caudate; NoRep, no reputation risk; Rep, reputation risk; BOLD, blood-oxygen-level-dependent response

A second GLM analysis (GLM2) allowed us to further investigate spontaneous lying under reputation risk, highlighting the differences between self-gain and other-gain motivations: i.e., SG-Lies and OG-Truths (unfavorable outcome); SG-Truths and OG-Lies (favorable outcome). We here focused on decisions made when the outcome was unfavorable because this outcome is arguably the most interesting condition as it contrasts SG-Lie and OG-Truth, which we are most interested in.

GLM2 results showed increased activity in the right ACC for SG-Lies compared to OG-Truths, regardless of the reputation condition (Fig. 2a, red region and Fig. 2b, red bars 1 & 4 vs. red bars 2 & 5 and Table 2a). Furthermore, an interaction effect for SG-Lies vs. OG-Truths with reputation risk was found in the bilateral ACC and in the left vmAI (Fig. 2a, blue regions and Fig. 2b, blue bar plots and Table 2b). This means that these regions are selectively more active for SG-Lies vs. OG-Truths under reputation risk as compared to no risk conditions (compare bar 4 & 5 vs. bars 1 & 2 in Fig. 2b). As these regions are elicited the most for SG-Lies under reputation risk (i.e., the main aim of the current study), we used them as our main regions of interest (ROIs) for the following correlation and connectivity analyses.

**Table 2.**
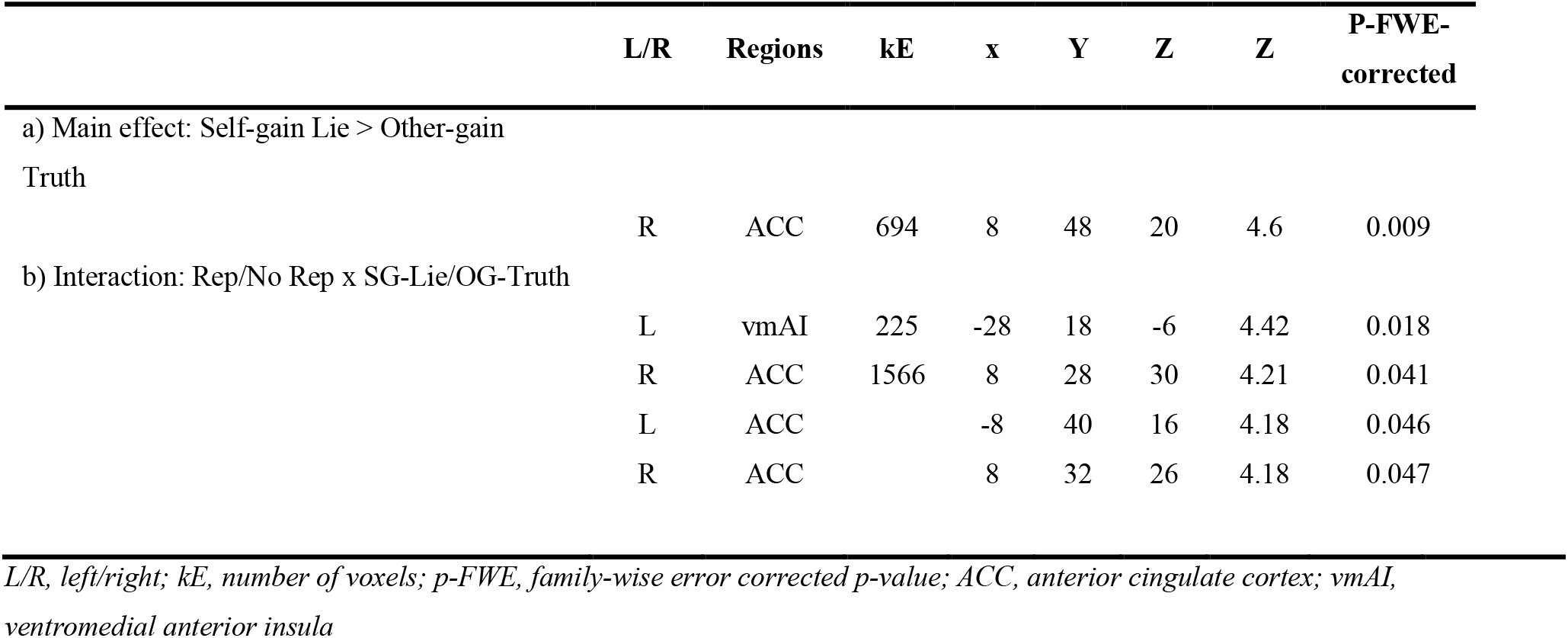
Brain activation in the contrast of, 1a) NoRep_SG-Lie + Rep_SG-Lie > NoRep_OG-Truth + Rep_OG-Truth, 1b) Rep_SG-Lie + NoRep_OG-Truth > NoRep_SG-Lie + Rep_OG-Truth. Only clusters and their local maxima that survived family-wise error correction (p <.05) are presented.

|  | L/R | Regions | kE | x | Y | Z | Z | P-FWE-corrected |
| --- | --- | --- | --- | --- | --- | --- | --- | --- |
| a) Main effect: Self-gain Lie > Other-gain Truth |  |  |  |  |  |  |  |  |
|  | R | ACC | 694 | 8 | 48 | 20 | 4.6 | 0.009 |
| b) Interaction: Rep/No Rep x SG-Lie/OG-Truth |  |  |  |  |  |  |  |  |
|  | L | vmAI | 225 | -28 | 18 | -6 | 4.42 | 0.018 |
|  | R | ACC | 1566 | 8 | 28 | 30 | 4.21 | 0.041 |
|  | L | ACC |  | -8 | 40 | 16 | 4.18 | 0.046 |
|  | R | ACC |  | 8 | 32 | 26 | 4.18 | 0.047 |

*L/R, left/right; kE, number of voxels; p-FWE, family-wise error corrected p-value; ACC, anterior cingulate cortex; vmAI, ventromedial anterior insula*

**Figure 2.**
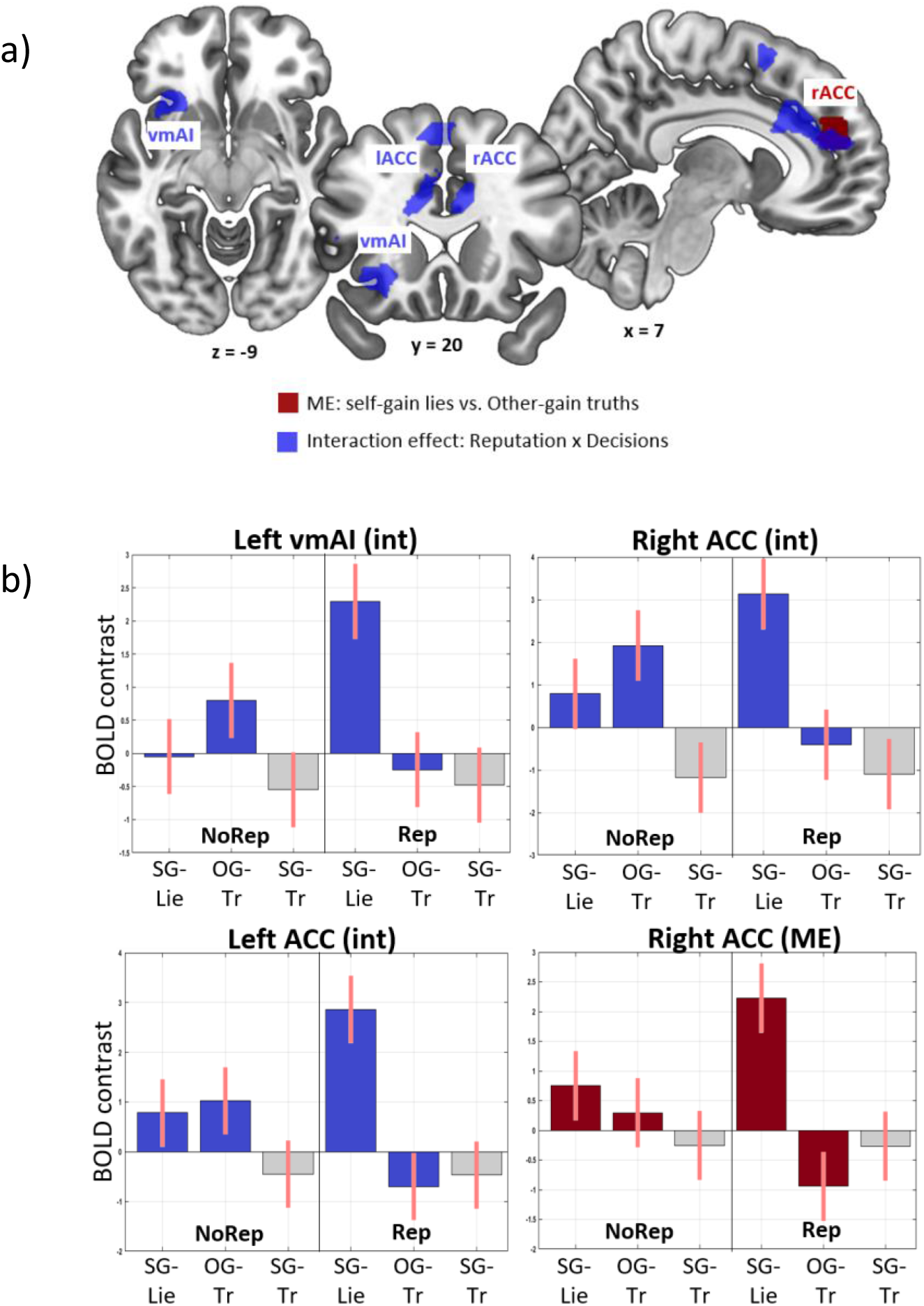
Results of GLM2: a) axial, coronal, and sagittal slices showing, in red, the regions active for self-gain lies vs. other-gain truths in both conditions and, in blue, the regions that are most active for self-gain lies vs. other-gain truths with reputation risk, which are left vmAI, and bilateral ACC (interaction effect: self-gain lies/other-gain truths x reputation/no reputation). b) Bar plots showing, in blue, the increased activation of the left vmAI and left and right ACC for self-gain lies with reputation risk (bar 4) and, in red, showing increased activation for self-gain lies (bars 1&4) vs. other-gain truths (bars 2&5) in both conditions. Abbreviations: vmAI, ventromedial anterior insula, lACC/rACC, left/right anterior cingulate cortex; ME, main effect; int, interaction effect; SG-Lie, self-gain lies; OG-Tr, other-gain truth; SG-Tr, self-gain truth.

The main effect of lies (self-gain) vs. truths (self-gain and other-gain) elicited similar regions as the main effect of GLM1 (see Fig. S2A, green map, and Table S4.1a) and the same interaction was found in the left ventromedial insula with reputation risk (see Fig. S2A (blue) and Table S4.1b). When contrasting SG-Lies with SG-Truths for both conditions bilateral ACC and AI activation is found, together with right SMA (see Fig. S2B, yellow map, and Table S4.2a). Finally, contrasting OG-Truths with SG-Truths for both conditions elicits bilateral ACC and right SMA (see Fig. S2.C, orange map, and Table S4.3a). No interaction effect with reputation was found for the final two contrasts.

### Behavioral results

We tested whether participants’ lying behavior (both for self-gain and other-gain motivations) was influenced by the fact that they were risking being caught by their opponent. Firstly, we tested whether the outcome (Fav or UnFav) and risk (No Rep or Rep) influenced lying percentages and response times. Secondly, we wanted to know whether certain dispositional personality factors predicted the probability of lying for both the no reputation risk and reputation risk condition. To deal with this question, we correlated questionnaire scores with other-gain (OG-Lies) and self-gain lying (SG-Lies) percentages in both conditions.

In agreement with previous literature^26^, the results of the binomial 2 × 2 generalized linear mixed model showed that lying percentage is modulated by outcome and reputation (Fig. 3a, Interaction effect: Lie ∼ Out x Rep: χ^2^(1) = 7.49, p = .006). As expected, when the outcome was unfavorable, participants lied significantly more for self-gain when they were guaranteed anonymity compared to when their reputation was at risk (z-ratio = 4.55, p < .0001). Moreover, fewer OG-Lies were made in the reputation condition compared to the no reputation condition (z-ratio = 3.14, p = .009). OG-Lies, however, were made much less often compared to SG-Lies in both conditions (No Rep: z-ratio = -6.16, p < .0003, Rep: z-ratio = - 5.82, p < .0001). Moreover, we found that manipulativeness scores correlated positively with SG-Lie percentage during reputation risk (Fig. 3b, r = .46, p = .01), while this was not significant for the no reputation condition (r = 0.29, p = 0.13). Thus, the more subjects were manipulative the more they tended to make SG-Lies in the reputation condition, while this positive correlation was not significant when they were anonymous. No other correlations were found between the lie probability and the BIDR and the CDM.

**Figure 3.**
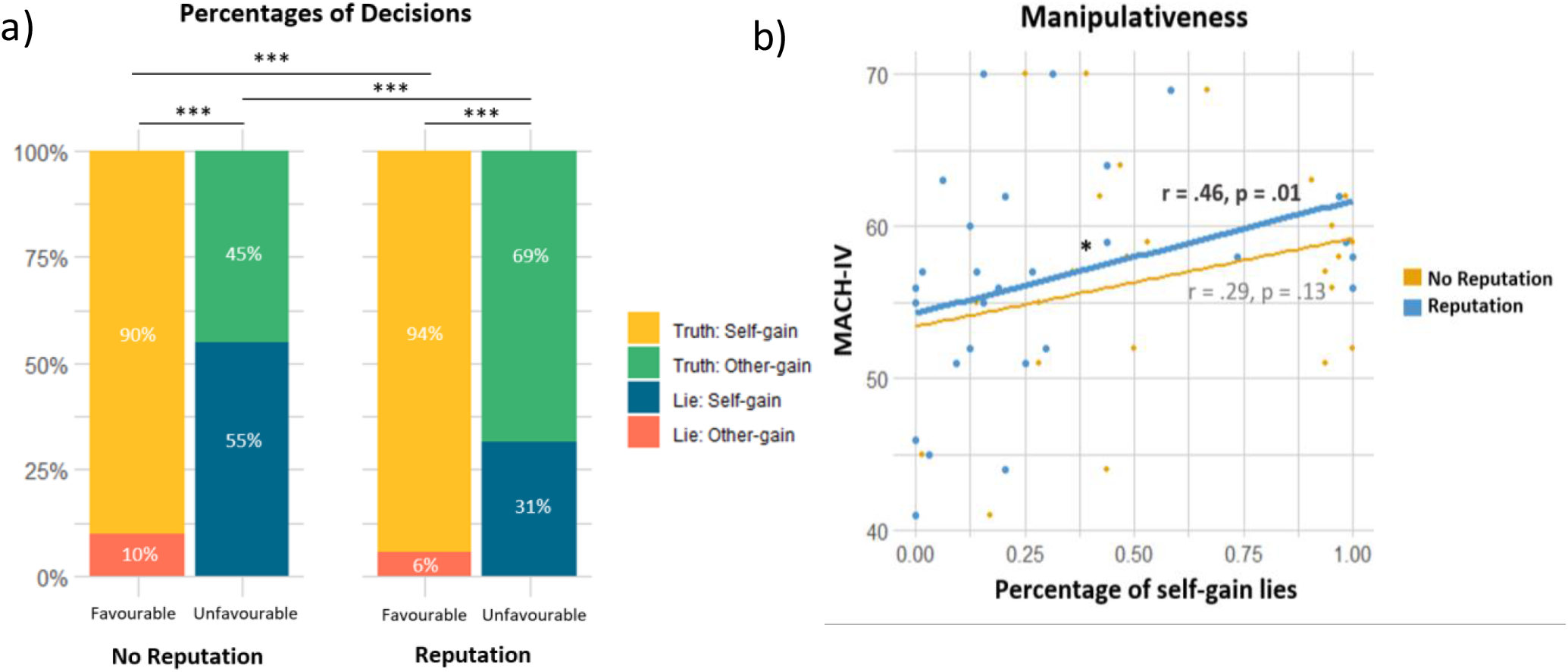
a) Percentages of decisions for each reputation condition (No Rep or Rep) and within each outcome (Fav or UnFav). b) Correlation graph of individuals Machiavellian scores with the percentages of self-gain lies during no reputation risk (yellow) and reputation risk (blue). The line plot in bold depicts significance. MACH-IV, Machiavellian questionnaire scores

Lastly, we find that truthful responses take significantly less time than lies [χ^2^(1) = 20.69, p < .0001] and decisions are made faster when the outcome is favorable [χ^2^(1) = 19.14, p < .0001] (see Fig. S3). A significant interaction effect was found between all 3 factors (reputation x outcome x decision: χ^2^(1) = 5.07, p = .02). Post-hoc tests revealed that for both the no reputation and the reputation condition, self-gain truths take the least amount of time compared to all the other conditions.

### Correlation analyses

We tested whether the BOLD signal of the main regions of interest found in the interaction effect (i.e., the left vmAI, and the left and right ACC) covariated with qualitative measures. We found an inverse relationship between MACH-IV scores and BOLD activity in the left ACC during self-gain lies under risk, i.e., the higher participants scored on manipulativeness the lower the activity in the left ACC for self-gain lies under the reputation risk condition (Fig. 4a, r = .54, p = .04). Contrarily, the left ACC was less active during other-gain truth under risk for less manipulative participants (Fig. 4b, r = -.66, p = .007). Less manipulativeness was also associated with increased activity in the left vmAI during NoRep_SG-Lies (see Fig. S4, r = -.60, p = 01).

**Figure 4.**
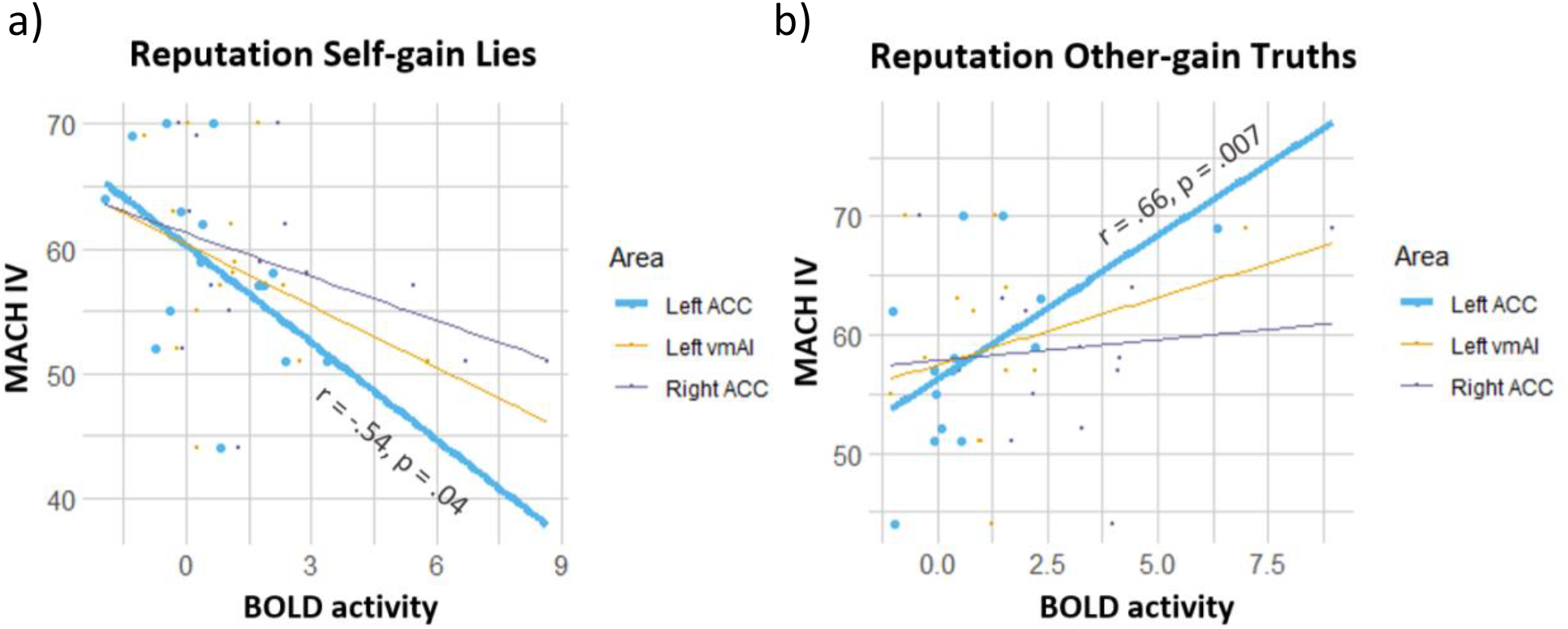
a) Correlation of averaged beta-values in 3 brain areas (LACC, LvmAI, RACC) during self-gain lies with reputation risk and Machiavellian scores, b) Correlation of averaged beta-values in 3 brain areas (LACC, LvmAI, RACC) during other-gain truths with reputation risk and Machiavellian scores. The line plots in bold depicts significance. Abbreviations: ACC, anterior cingulate cortex; vmAI, ventromedial anterior insula

### Connectivity analyses

Finally, we explored whether and how the pattern of functional connectivity changed for the three ROIs derived from the interaction effect (i.e., the left vmAI, and the left and right ACC) of SG-Lies vs. OG-Truths as a function of reputation risk. Relative to other-gain truth-telling, self-gain lying with reputation risk induced a stronger coupling of the left vmAI with the bilateral ACC: right ACC: F(2,13) = 7.59, p = .006, left ACC: F(2,13) = 4.82, p = .02 (Fig. 5a & b, yellow bar vs. purple bar). In the no reputation risk condition, this relationship is reversed (see Fig. 5a & b, red bar vs. blue bar). The same contrast showed, however, strongest connectivity between the right ACC and the left dmPFC for Rep_SG-Lie [F(2,13) = 4.22, p = .03] (Fig. 5c, yellow bar).

**Figure 5.**
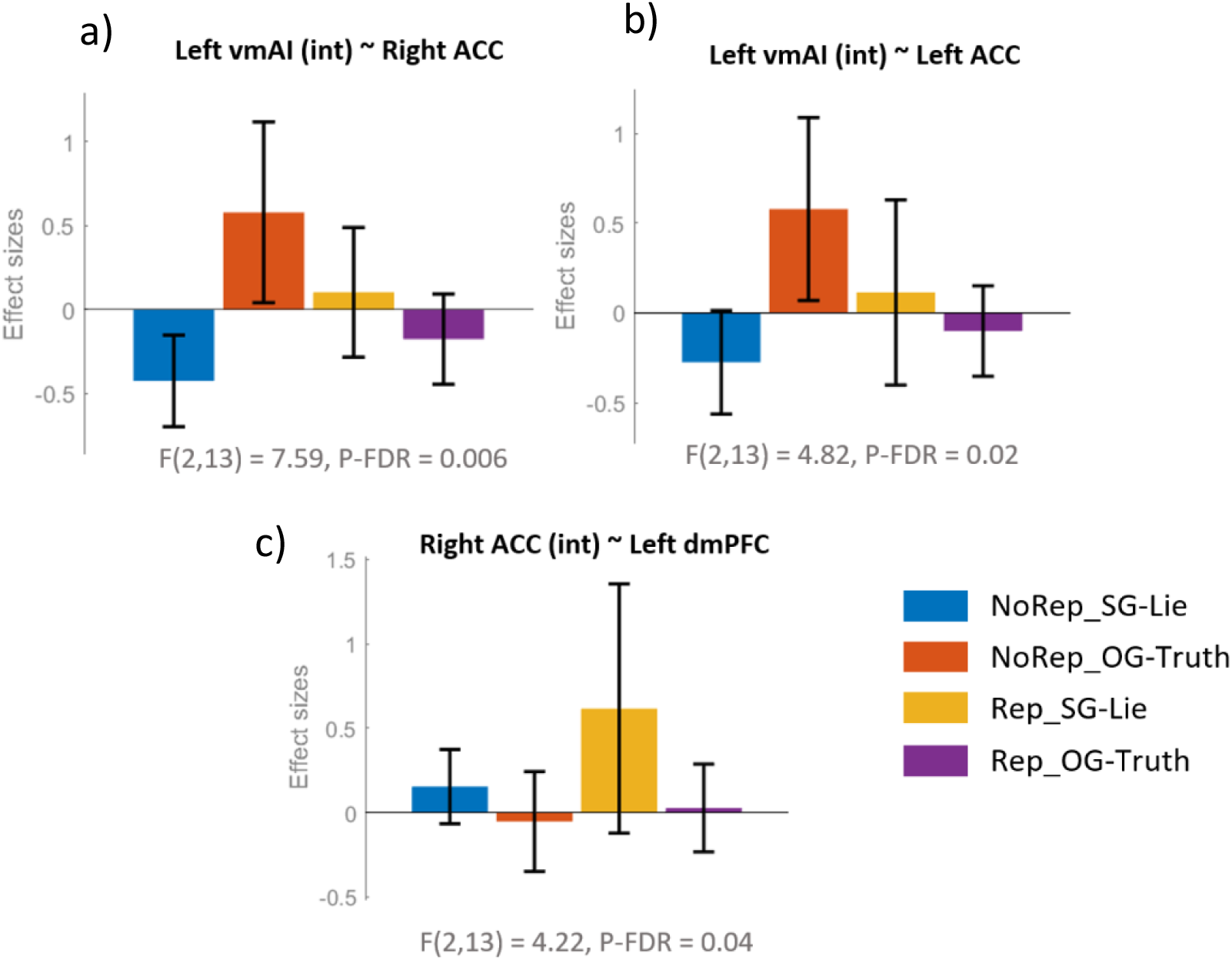
Bar plots of connectivity analysis. a) stronger connectivity of left vmAI and right ACC for self-gain lies with reputation vs no reputation risks (yellow vs. blue) and for other-gain truths with no reputation vs. reputation risk (red vs. purple), b) stronger connectivity of left vmAI and left ACC for self-gain lies with reputation vs no reputation risks (yellow vs. blue) and for other-gain truths with no reputation vs. reputation risk (red vs. purple), c) stronger connectivity of the right ACC and left dmPFC for self-gain lies with reputation risk vs. other conditions (yellow vs. blue, red, purple). Abbreviations: vmAI, ventromedial anterior insula; ACC, anterior cingulate cortex; dmPFC, dorsomedial prefrontal cortex; NoRep_SG-Lie, self-gain lies without reputation risk; NoRep_OG-Truth, other-gain truths without reputation risk; Rep_SG-Lie, self-gain lies with reputation risk; Rep_OG-Truth, other-gain truths with reputation risk; P-FDR, false discovery rate corrected p-value.

## Discussion

Using an ecological approach wherein participants could freely decide to lie or to tell the truth we examined the modulation of neural activity that reputation risk and game outcome factors exerted on spontaneous dishonest social decision-making. We have been able to determine the regions more active when spontaneous lies vs truths were made independently from reputation condition and motivation of decisions (self-gain or other-gain). Notably, we highlighted the brain areas recruited more for lies in general and those activated by self-gain lies in reputation risk conditions. Finally, we have been able to determine the strength of the connections between the nodes in the neural circuit underpinning the deceptive and truthful decisions in our ecological task.

### Brain regions involved in lying compared to truth-telling

When exploring the brain regions associated with lying relative to truth-telling regardless of the reputation risk and outcome factors, we found changes of activity in a prefrontal circuit, including the bilateral ACC and AI, the left dlPFC, the SMA, and the right caudate. The activation of these anterior regions of the brain is in good agreement with meta-analyses on the neural correlates of dishonesty^18–20,32^. In particular, activation in the ACC and the insula have been consistently found for deceptive compared to truthful behavior across a variety of tasks and stimuli. The ACC has been linked to a wide range of executive control processes that are needed to execute a deceptive response, such as social context integration^33^, working memory, inhibition of competing responses, mediation of cognitive conflict^34,35^, task-switching, reward, interoception and motivation in decision-making^2,20,36–39^. The anterior insula, besides its general involvement in executive control, has also been linked to the visceral responses (e.g., blood pressure, heart rate, body temperature) and interoceptive activity^40^ that accompany deceptive behavior^19,20^. This adds to the notion that AI may be an integrative hub that detects bottom-up salient events, then projects information to task and context-relevant brain networks, to access attention and working memory resources^41^ and to make decisions operational^42,43^. Moreover, the dlPFC has been linked to spontaneous vs. instructed dishonesty^44^ and has been found to play a role in executive processes regardless of the deception task used^45^. Lie-related activation of the right caudate might be related to the inhibition of socially accepted responses, in this case, telling the truth^19^ or anticipation and valuation of reward. It is worth mentioning that higher resting-state connectivity of reward processing with the nucleus caudate as central node was directly correlated with propensity to cheat ^15^. Moreover, resting-state connectivity of ROI’s for reward, self-reference, morality and cognitive control were found to be predictive of dishonesty rate ^46^. Overall, the activations found for spontaneous deception in our study are consistent with those found in the previous literature, indicating that the paradigm effectively promotes deception.

### Brain regions involved in self-gain lying when reputation is at risk

Our ecological task allowed us to explore the activity of lying-related network of regions when the risk of being caught in a lie was or was not present. Under reputation risk, we found a selective activation of the ventromedial portion of the left anterior insula for self-gain lies. The anterior insula is known not only for its role in interoceptive awareness and visceral responses^47^, but also for its involvement in socially relevant functions like social exclusion^48^, exposure to unfair treatment^49^, anticipation of reputation decisions^42^, and when making inequitable decisions^50^. Studies have shown that the activation in the AI is predictive of subsequent immoral behavior^51^ and its activation is associated with anticipation of guilt, and emotion that is crucially involved in moral decision making^52,53^. Lying for altruistic goals compared to self-serving goals has been found to reduce AI activity^54^. Research in neuroeconomics has shown that the processing of financial risk-taking when there are potential losses involved is mainly represented in the anterior insula^49,55,56^. It is hypothesized that this is due to the processing of more aversive emotions related to risk-taking or risk anticipation^57^ than control conditions. Specifically the ventral AI, compared to the dorsal AI, has been associated with more affective processes, e.g., mediating aversive feelings that generate motivation to norm enforcement^58^. Based on these findings, a plausible interpretation of the activation of the left vmAI during deception with reputation risk found in our study is likely due to heightened emotional processing, specifically due to aversive emotions that arise with risk-taking, such as fear, sadness, disgust, anxiety^59,60^or guilt^61^. Here we further add to the literature that the left vmAI plays a role in the modulation of dishonest decision-making under reputation risk.

Diving deeper into the neural nature of decision-making under reputation risk, we found that when the outcome was unfavorable, i.e., the opponent wins and the participant’s tendency to make self-gain lies is maximal, there is an increase in vmAI activity together with the left and right ACC for dishonesty with a reputation risk. Interestingly, that the greater recruitment of the ACC during deception in a social context compared to non-social deception studies may indicate greater conflict monitoring as individuals are, under social rules, supposed to act honestly in social vs. non-social context and therefore increased executive control processes are needed to control the ‘usual’ honest response^19^. It has also been reported that the presence of an audience reduced the likelihood of accepting an immoral offer for monetary gain and that audience vs. no audience engaged a brain network including the anterior insula, the ACC, and the right TPJ^63^ and that these regions may reflect meta-representations of what other people think about us or of our desire for social norm compliance^58^. Going against social norms has also been argued to have potentially higher emotional costs^62,63^. In incarcerated psychopaths, a reduces ACC activity was found when making dishonest decisions, arguably due to their lack of moral concern ^34^.

Based on these studies, we argue that going against social norms (in this case, honesty) in favor of a monetary reward, while there is a risk of getting caught, is emotionally more salient than going against social norms without social risk. This may entail increased bottom-up activity in the vmAI and, in turn, the recruitment of the ACC needed for increased executive control and mediation of cognitive conflict. Interestingly, this is in line with a study in which we showed that to produce a self-gain lie when reputation is at risk, people need to regulate their sympathetic activation more, as indexed by higher temperature on the nose^25^.

Our findings are in seeming contrast with a study reporting increased subgenual ACC activity during deceptive decisions when there was no risk of confrontation^64^. However, risk of confrontation in this study means monetary penalization, while in ours it means loss of reputation without loss of money. This difference could make one task more sensitive than the other in grasping the role of the ACC in monitoring reputation risk.

It is also noteworthy that our functional connectivity analysis showed a stronger coupling of the left vmAI with the left and the right ACC during self-gain lies compared to other-gain truths in the reputation risk condition. Tellingly, this relationship was reversed in the no reputation condition. This pattern of results may reflect the interplay between emotional and executive processes. Cognitive conflict seems to arise both when selfish lies are made with a potential social penalty but monetary reward and when altruistic truths are told with no social reward and a monetary penalty. The increased connectivity of the ACC and the left dlPFC during dishonesty was also found in a study in which bottom-up detection of conflict involves the recruitment of the ACC, that, in turn, recruits the dlPFC for top-down self-regulatory processes^34^. One key addition of our study is that this connectivity pattern becomes stronger when reputation is at risk. Moreover, in keeping with previous studies, the pattern of connectivity reported in our study might indicate a need for top-down control associated with conflict-induced behavioral adjustments^65^ and motor-control functions^66^.

### Individual differences in behavioral and brain correlates of deception

Another interesting result of our study is that manipulative characteristics of our participants correlated both with the tendency to deceive and the brain correlates associated with this behavior. Higher Machiavellian scores were associated with a higher production of self-gain lies in the risk reputation condition and with a decrement of activity in the left ACC in the very same condition. Individuals with higher MACH-IV scores typically have less problems deviating from social norms. That more manipulative individuals making risky selfish lies do not need to recruit the ACC as much may suggest that their decision to violate the norm requires less conflict monitoring and therefore less executive control. Both findings are in accordance with our previous research indicating that higher manipulative traits were associated with a smaller effect of reputation risk^26^, a smaller inhibition of the cortical motor readiness to lie^27^ and a smaller regulation of the sympathetic system during lying when reputation is at risk^25^.

In the present study, we also found reduced left ACC activity for other-gain truth-telling under reputation risk in less manipulative individuals. In sum, high manipulative individuals need strong cognitive control when making honest decisions under risk and low manipulative individuals need it when making dishonest decisions under risk. These results are in keeping with the theory^21^ that cognitive control is needed to override one’s own moral default^16^.

### Behavioral findings

We found that both reputation risk and outcome factors influenced participants’ dishonest decisions, in the sense that, risking one’s own reputation reduces self-gain dishonesty and unfavorable game outcome increases it. These behavioral findings are in line with those reported by Panasiti and colleagues^25–27^ and Azevedo and colleagues^28^. The finding that other-gain lies are higher in the no reputation risk may be linked to the human tendency of rather donating anonymously than with a chance of being found out^68^. That this result was not found in a previous study where we used a similar protocol^26^ may be due to the difference in effect size of the reputation condition on lying behavior. Moreover, we found that lying takes longer than truth-telling corroborating previous findings ^35,54,69^. However, there was no significant difference in reaction times between outcome (unfavorable vs favorable) and reputation risk conditions.

### Possible limitations and conclusions

Unlike what typically happens in classical experimental designs where the same number of observations is obtained for each condition, we had to deal with a somewhat unbalanced data set since, given the ecological nature of the task, the different participants made rather diverse choices. We acknowledge that, on the one hand, this can be considered as a limitation. On the other hand, however, our approach provides a veridical picture of what happens under life circumstances where complex decision behaviors come with high interindividual variability at both dispositional and situational levels.

In conclusion, our experimental approach allowed us to reveal some of the important regions needed for making dishonest decisions when one’s own reputation was or was not at risk. The recruitment and the increased functional connectivity of the anterior insular and cingulate cortex when making dishonest decisions under the influence of a social risk points towards a greater need of emotional processing and executive functioning, likely because going against one’s own social norms (honesty before money) causes inner conflicts. When one’s own social norms, however, shift toward selfishness (money before honesty) like in the more manipulative individuals, the need for conflict monitoring seems to arise more with altruistic honest decisions and less with selfish dishonesty decisions in social context.

## Materials and methods

### Participants

Thirty-four participants enrolled in the fMRI experiment. All participants had normal or corrected-to-normal vision, were free from any contraindication to fMRI, and had no history of major psychiatric or neurological problems. All participants gave their written informed consent, and the study was approved by the independent Ethics Committee of the Santa Lucia Foundation IRCCS (Scientific Institute for Research Hospitalization and Health Care). Data from six participants were excluded prior to analyses (three participants were excluded due to technical problems related to the acquisition of the anatomical scan; two did not believe that the opponent was a real player; and one did not understand the task properly), leaving a final sample of 28 participants (range = 20-45 years; mean = 26 years, SD = 6 years).

The appropriate sample size for this study was estimated with G*Power 3.1.9.2 (ANOVA, repeated measures, within factors), considering a medium effect size of 0.20 (predicted based on Panasiti et al., 2011^26^, using the same design), a significance level of 0.05, 1 group, 12 measurements (i.e., 3 reputation condition x 2 decisions x 2 outcomes). This indicated a power > 95% using a sample size of 28 participants.

### Task

We used the TLCG, a card game to explore dishonest decision-making in a social context. This type of paradigm proved adept to highlight situational and dispositional factors as well as social variables that may influence the participants’ performance ^25–28^. The TLCG was adapted for the scanner session. Inside the MRI scanner, participants played the TLCG against a computerized opponent (OP). Crucially, however, participants were told they were playing against a real opponent who was sitting in a different room and whom they would meet at the end of the experiment. Only at the end of the experiment were participants fully debriefed and told that the OP’s choices were made by a computer algorithm.

Each trial of the TLCG started with the OP choosing one of two covered cards, one on the left and one on the right side of the display, within a time window of 1 – 2.5 s (Fig. 6a). The chosen card could be either the ace of spades or the ace of hearts, indicating a loss for the OP (favorable outcome for participant) or a win for the OP (unfavorable outcome for participant), respectively. Prior to the experiment, participants were informed that only they could see the outcome of the OP’s choice and communicate it to the OP. Participants were free to communicate the outcome truthfully or to reverse the outcome. Decisions were communicated by pressing one of two response keys within a time window of 2.5s. For instance, the participant could communicate that the OP chose the losing card while, in reality, the OP had chosen the winning card (i.e., Self-gain lie) or s/he could communicate that the OP chose the winning card while OP had chosen the losing card (i.e., Other-gain lie). After a jittered inter-trial interval (mean of 1.75 s, range = 1 - 2.5 s) filled with a fixation cross, a new trial began. Participants were aware that in each trial the monetary pay-off could go only to the winner and that a different amount of money was associated with each trial. The exact gain would only be communicated at the end of the game to rule out that participants choice was based on a trial-by-trial computation of gain and loss, and to ensure that the temptation to deceive was comparable across trials. Participants were given 10 euros for their participation and could win up to 25 euros extra during the game.

**Figure 6.**
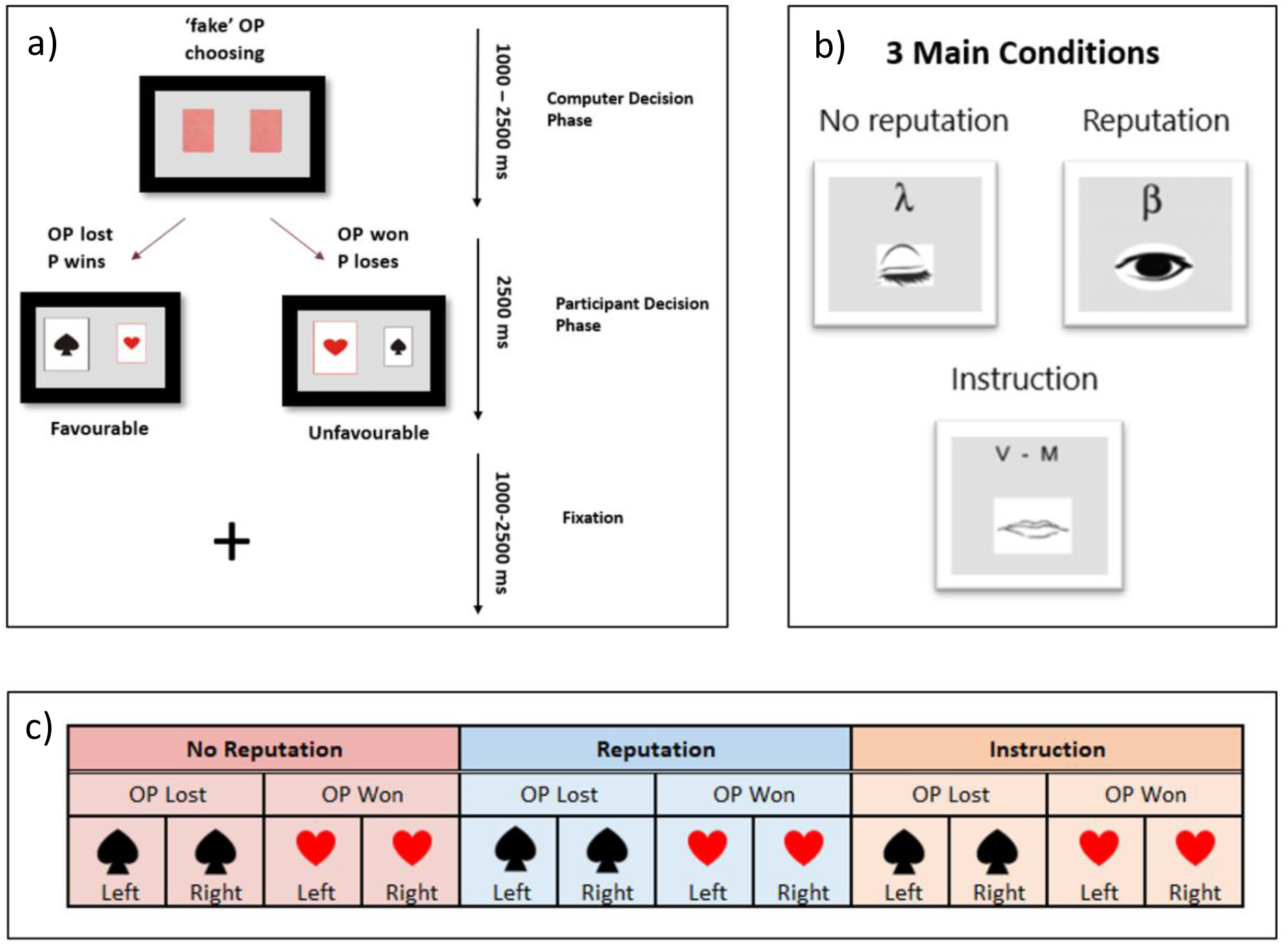
a) Time-course of a representative trial, starting with the opponent (OP) choosing one of two covered cards (1000-2500 ms; computer decision phase) ending with the exposure of the selected card to the participant. In the participant decision phase (in a time-window of 2500 ms), they had to decide what to communicate back to OP by either lying and thus changing the outcome (selecting the not-chosen card) or telling the truth and thus respecting the outcome (selecting the chosen card). The trial ended with a fixation point in a jittered inter-trial interval (1000-2500 ms). b) The 3 main conditions and their symbols during the experiment. c) Visual representation of 3 × 2 × 2 experimental design: experimental condition (No rep, Rep, Ins) x the possible outcome (OP lost (Fav) or OP won (UnFav)) x position of selected card by OP (left or right)

The game was performed under 3 main conditions (Fig. 6b), two spontaneous and one instructed: (1) No reputation risk (No Rep; indicated with an eye closed symbol and a λ, lambda), in which the participants knew their decision would be unknown to the OP; (2) Reputation risk (Rep; indicated with an eye open symbol and a β, beta), in which they knew there was a 75% chance of the OP finding out their decision; and (3) Instruction (Ins; indicated with a mouth symbol and either the letter “V” for the Italian word “Verità”, truth, or the letter “M” for the Italian word “Menzogna”, lie, depending on whether participants were instructed to lie or telling the truth). In this condition, participants were instructed about which specific decision they had to make. The experiment included a total of 384 trials, given by the crossing of 2 (left vs. right card selected by OP) by 2 (favorable vs. unfavorable outcome for participants, i.e., OP chose ace of spades or ace of hearts, respectively) by 3 (No Rep, Rep, or Ins experimental conditions) by 32 repetitions of each type of trial. The total experiment consisted of 4 functional MRI runs including 96 trials each. These were administered in 9 mini-blocks, 3 for each condition (i.e., No rep, Rep, or Ins), with different lengths, namely, either 8, 10 or 14 trials, to avoid predictability of the number of consecutive trials. Participants received no feedback about whether they were caught lying during the functional scans. Moreover, to avoid outcome predictability, the four types of possible outcomes (Fig. 6c) were randomized across the whole experiment, and not within each mini-block. The participant could make 4 types of decisions depending on the outcome: OP won, i.e., unfavorable outcome (UnFav), eliciting either a self-gain lie (SG-Lie) or an other-gain truth (OG-Truth), and OP lost, i.e., favorable outcome (Fav), eliciting either an other-gain lie (OG-Lie) or a self-gain truth (SG-Truth). Instruction trials were used as control conditions in fMRI analyses.

### Functional Magnetic Resonance Imaging

All images were acquired with a Siemens Allegra fMRI scanner (Siemens Medical Systems, Erlangen, Germany) operating at 3T. A quadrature volume head coil was used for radio-frequency transmission and reception. Head movements were minimized by mild restraint and cushioning. For each subject, functional MR images were acquired using echo-planar imaging (EPI) [slices = 32, TR = 2.08 s, TE = 30 ms, in-plane resolution = 3 × 3 mm2, slice thickness = 2.5 mm, flip angle = 70 degrees), covering the entire cortex. Structural MR images were obtained using a T1-weighted 3D magnetization prepared rapid gradient echo (MPRAGE) imaging sequence [slices = 176, TR = 2s, TE = 4.38 ms, in-plane resolution = 0.5 × 0.5 mm2, slice thickness = 1mm, flip angle = 8 degrees]. For each participant, we acquired 1284 fMRI volumes, 321 for each of the four functional runs. The first four volumes of each run were used for stabilizing longitudinal magnetization and then discarded from further analysis.

### fMRI data analysis

#### Pre-processing

The fMRI data were pre-processed and analyzed with the Statistical Parametric Mapping package SPM12 (www.fil.ion.ucl.ac.uk) implemented in MATLAB R2019b (The MathWorks, Natick, MA). Using the ARTrepair toolbox (https://www.nitrc.org/projects/art_repair/), all images were previewed (art_movie) for detection of excessive motion artifacts and bad slices were detected and repaired (art_slice) by interpolating adjacent slices. Functional images were then slice-time corrected to compensate for slice acquisition delays between the first and last slice by using the middle slice as a reference. Subsequently, images were realigned and unwarped to correct for head movement. Images were registered to the first volume using a 2nd degree B-spline interpolation, whereas during the unwarp re-slicing a 4th degree B-spline interpolation was used. Individual structural T1-weighted images were segmented into grey matter (GM), white matter (WM) and cerebrospinal fluid (CSF) using SPM tissue probability maps. Structural images were bias-corrected with a light regularization and a 60 mm cut-off, while forward deformation fields were created. The segmented structural bias-corrected image was then co-registered with the slice-timed, realigned, and unwarped functional images, using the mean unwarped image as the source image. The obtained forward deformation fields during segmentation were used during normalization to bring co-registered functional images of the 4 sessions into the MNI space (2 mm isotropic voxel) using the 4th-degree B-Spline interpolation. Finally, images were smoothed with a Gaussian kernel of 8 mm FWHM to ameliorate differences in inter-subject localization.

The pre-processed images were then examined again using the Artifact Detection Tool (2015) software package (https://www.nitrc.org/projects/artifact_detect), for detecting those scans in which – notwithstanding the above “repair” procedure – the excessive motion remained. Outlier scans were identified in the temporal differences series by assessing between-scan differences (Z-threshold: 3.0 mm, scan-to-scan movement threshold: 1 mm; rotation threshold: 0.02 radians). The outlier scans (3.1% overall) were omitted from the analysis by including a single regressor for each outlier in the design matrix.

#### fMRI General Linear Model analyses

The main aim of this study was to investigate how one’s decision to deceive modulates neural activity under the risk of losing one’s reputation. For this purpose, two separate general linear models (GLM1 and GLM2) were generated. GLM1 was performed to look at the overall neural difference between spontaneous lying and truth-telling, with an emphasis on how including a reputation risk factor modulates this difference, irrespective of the nature of the decision (self-gain or other-gain). GLM2 provides a more detailed picture of the regions involved in self-gain vs. other-gain decisions.

For each GLM, the statistical inference was based on a random-effects approach comprising two steps: first-level multiple regression models estimating contrasts of interest for each participant, followed by the second-level analyses for statistical inference at the group level. Non-sphericity correction^70^ was applied to account for possible differences in error variance across conditions, arising - for example - because of the different number of trials in the conditions of interest and/or any non-independent error terms for the repeated measures (see second-level analyses).

Similar to previous studies on spontaneous deception ^5,18,69,71,72^, the naturalistic nature of the task caused an imbalance in the number of trials for both the reputation and no reputation conditions; e.g., participants who always chose other-gain truths instead of self-gain lies or vice versa. This meant that some participants had to be excluded due to an insufficient number of trials. In our first GLM, both types of lies (other-gain and self-gain) and both types of truths (other-gain and self-gain) were collapsed to contrast overall lying with truth-telling, meaning that, for example, insufficient trials for other-gain lies would be compensated with sufficient trials for self-gain lies. However, in our second GLM, all specific decisions were regressed separately meaning more imbalance and therefore, higher rates of participant exclusion.

For GLM1, data from 6 participants were excluded from the analyses due to insufficient trials, based on a criterion of less than 10% for lying or truth-telling decisions per condition (No Rep or Rep) per outcome (Fav or UnFav), i.e., 64 trials per condition meaning less than 6.4 trials (see Supp. Mat. Table S1 for lie/truth count per condition for each participant). This left twenty-two participants (mean age = 25.4, SD = 4.1, range = 20 – 32 years) in GLM1.

Six regressors of interest were modelled at the first level, corresponding to the following conditions: 1) spontaneous lies with no reputation risk (NoRep_Lie); 2) spontaneous truths with no reputation risk (NoRep_Truth); 3) spontaneous lies with reputation risk (Rep_Lie); 4) spontaneous truths with reputation risk (Rep_Truth); 5) instructed lies (Ins_Lie); 6) instructed truths (Ins_Truth) (see Supp. Mat. Fig. S1 for a visual representation).

Additionally, six sets of motion parameters derived from the realignment stage and outlier regressors were included as covariates of no interest. The events were modelled as mini blocks, time-locked to the onset of the decision phase, with a duration equal to the time window of the decision phase (i.e., 2.5 s). All regressors were convolved with the canonical haemodynamic response function (HRF), and a temporal high-pass filter with a cut-off at 128s was applied to reduce low-frequency noise. For each participant, linear contrasts were used to average the parameter estimates associated with each of the six conditions of interest, across the four fMRI-runs. For the group-level analysis, we carried out a within-subject ANOVA with factors: Condition (no rep, rep, ins) and Decision (lie, truth).

For GLM2, data from 13 participants were excluded based on the 10% criterion per decision (Self-gain Lie, Other-gain Truth, Self-Gain Truth) within each condition (No Rep or Rep) and Outcome (Fav or Unfav) (see Supp. Mat. Table S2). Insufficient trials of other-gain lies (OG-Lie) were not considered since very few participants chose to lie for other-gain reasons (5.7% of NoRep trials and 5% of Rep trials across participants). For this reason, the other-gain lies were modelled at first level but not included in the second level model. GLM2 included a total of fifteen participants (mean age = 25.2, SD = 4.16, range = 20 – 32 years). Twelve regressors were modelled at first-level: 1) self-gain lies with no reputation risk (NoRep_SG-Lie); 2) other-gain truths with no reputation risk (NoRep_OG-Truth); 3) other-gain lie with no reputation risk (NoRep_OG-Lie); 4) self-gain truth with no reputation risk (NoRep_SG-Truth); 5) self-gain lies with reputation risk (Rep_SG-Lie); 6) other-gain truths with reputation risk (Rep_OG-Truth); 7) other-gain lie with reputation risk (Rep_OG-Lie); 8) self-gain truth with reputation risk (Rep_SG-Truth); 9) instructed self-gain lies (Ins_SG-Lie); 10) instructed other-gain truths (Ins_OG-Truth); 11) instructed other-gain lie (Ins_OG-Lie); 12) instructed self-gain truth (Ins_OG-Truth) (see Supp. Mat. Fig. S1 for a visual representation).

As in GLM1, this model included motion parameters and outlier regressors as covariates of no interest; the events were modelled as mini-blocks time-locked to the onset of the decision phase (duration = 2.5 s); all regressors were convolved with the HRF, with a cut-off filter at 128s, and linear contrasts were used to average the parameter estimates associated with each of the twelve conditions of interest, across the four fMRI-runs. However, we did not include in the group-level analysis, the “other-gain lies” regressors for all the three conditions (NoRep, Rep and Ins) due to insufficient trial count across participants that hampers good parameter estimates. We, therefore, conducted another within-subjects ANOVA including Condition (NoRep, Rep, Ins) x Decision (SG-Lie, OG-Truth, SG-Truth).

Our contrasts of interest for GLM1 were the main effect of spontaneous lying vs. truth-telling (NoRep_Lie + Rep_Lie > NoRep_Truth + Rep_Truth) and the interaction effect of reputation and no reputation with lies and truths (Rep_Lie + NoRep_Truth > NoRep_Lie + Rep_Truth). For GLM2, we were specifically interested in clarifying the correlates of SG-Lies vs. OG-Truth for both Rep and NoRep conditions in unfavorable outcomes.

For both GLM1 and GLM2, in line with the main aim of the study, we constrained the search volume (using the small volume correction SPM function) within the brain areas responding to spontaneous decisions, i.e., all “spontaneous vs. instructed” trials, using a thresholded contrast image of p_uncorrected_ < .005. Data are presented including all significant activations at cluster-level using family-wise corrected p-values (significance at P_FWE-corrected_ < .05). Additionally, local maxima in the clusters were included when P_FWE-corrected_ < .05 at peak-level.

### Questionnaires

After the task and outside the fMRI scanner, the participants were qualitatively debriefed about their experience. A questionnaire was given right after the scanning session (See Supp. Mat. Q1). Two participants declared they did not believe that the OP was a real player and thus were excluded from the analysis. After this, the participants were administered: the Balanced Inventory of Desirable Responding (BIDR)^73^, which consists of two 20-item subscales, ranging from 20-140 and measures self-deception and impression management, both related to social desirability; the Machiavellianism Scale (MACH-IV)^74^, which is a 20-item scale where scores can range from 40 to 160 and measures the ability to use deception and manipulation to acquire power during everyday life interactions; and the Civic Moral Disengagement (CMD)^75^, a 40-item questionnaire, scoring from 40 to 200, and measures an individual’s tendency to make use of self-dismissal when violating civic duties and obligations in order to soften the moral consequences of their behavior ^76^. Questionnaires were chosen based on the study of Panasiti et al. (2011)^26^.

### Behavioral Data Analysis

To test whether Reputation (No Rep vs. Rep) and Outcome (Fav vs. UnFav) affected lying percentage, a 2 × 2 binomial generalised mixed linear model was used, with decision as the dependent variable (lie = 1 and truth = 0) and subject as a random factor. To analyze whether response times differed between Reputation (No Rep vs. Rep), Outcome (Fav vs. UnFav) and Decisions (lie vs. truth), a 2 × 2 × 2 general linear model was used with response time as the dependent variable and subject as a random factor. Type III Wald Anova function from the R package was used to determine the statistical significance of the fixed effects for both models. Least square means (from the *lsmeans* package) and Tukey corrections were used for post-hoc comparisons of the interaction effects. Finally, questionnaire scores were correlated with lying percentage scores for each spontaneous condition (No Rep and Rep) using Spearman’s correlation test. Only results passing a significance level of p < .05 are illustrated and included in the Result’s section.

### Correlation analyses

A Spearman’s correlation analysis was conducted to evaluate whether bold activity in the brain regions highlighted by GLM1 and GLM2 were correlated with questionnaire measures (i.e., MACH-IV, BIDR and CMD). Questionnaire scores were correlated with the bold signal extracted eigenvalues for each condition. For this, we used an 8 mm sphere (matching the Gaussian kernel) centered around all significant peaks that survived family-wise error correction during the GLM analyses. In the main text, we included correlations of regions of interest that are specifically linked to SG-Lies (GLM2) when reputation was at risk. All other correlations are available in the supplementary materials.

### Task-based connectivity analyses

Finally, we explored task-modulated functional connectivity to investigate potential connectivity differences in regions uncovered in GLM1 and GLM2 within the restricted search volume mentioned above. To this end, we used a generalized psycho-physical interaction (gPPI) approach implemented through the CONN toolbox (www.nitrc.org/projects/conn). For each participant, we constructed two different PPI–GLMs (one based on each GLM). Regions of interest (ROIs), defined based on our GLM analyses (see Table S3), were established using 8-mm spheres (matching the Gaussian kernel and built by the SPM toolbox MarsBaR; Brett et al., 2002) centered on the peak voxels from significant clusters of the contrasts (see Table 4.1a&b and Table S4.1a). For each ROI, bivariate regression matrices were calculated, yielding standardized regression coefficients that estimated the functional connectivity at the group level. Only results with FDR-corrected p-values < .05 are discussed.

## Author Contributions

Conceptualization, S.M.A., R.A. M.S.P.; Methodology and analysis, L.D., V.S., R.A. Writing – Original Draft, L.D., V.S., S.M.A; Writing – Review & Editing, all the authors; Funding Acquisition, S.M.A.; Resources, S.M.A.; Supervision S.M.A. and V.S

## Acknowledgments

The authors would like to thank Marco Tullio Liuzza and Emiliano Macaluso for their involvement in the preliminary stage of the study.

## Inclusion and diversity

We worked to ensure sex balance in the recruitment of human subjects and to prepare the study questionnaires in an inclusive way.

